# DNA methylation as a contributor to dysregulation of *STX6* and other frontotemporal lobar degeneration genetic risk-associated loci

**DOI:** 10.1101/2025.01.21.634065

**Authors:** Naiomi Rambarack, Katherine Fodder, Megha Murthy, Christina Toomey, Rohan de Silva, Peter Heutink, Jack Humphrey, Towfique Raj, Tammaryn Lashley, Conceição Bettencourt

**Affiliations:** Department of Neurodegenerative Disease, UCL Queen Square Institute of Neurology, London, UK; Department of Clinical and Movement Neurosciences, UCL Queen Square Institute of Neurology, London, UK; The Francis Crick Institute, London, UK; Reta Lila Weston Institute, UCL Queen Square Institute of Neurology, London, UK; German Center for Neurodegenerative Diseases, Tübingen, Germany; Nash Family Department of Neuroscience and Friedman Brain Institute, Icahn School of Medicine at Mount Sinai, New York, NY USA

**Keywords:** Frontotemporal lobar degeneration, Frontotemporal dementia, Progressive supranuclear palsy, DNA methylation, Epigenetics, Neurodegeneration, disease risk

## Abstract

Frontotemporal lobar degeneration (FTLD) represents a spectrum of clinically, genetically, and pathologically heterogeneous neurodegenerative disorders. The two major FTLD pathological subgroups are FTLD-TDP and FTLD-tau. While the majority of FTLD cases are sporadic, heterogeneity also exists within the familial cases, typically involving mutations in *MAPT, GRN* or *C9orf72,* which is not fully explained by known genetic mechanisms. We sought to address this gap by investigating the effect of epigenetic modifications, specifically DNA methylation variation, on genes associated with FTLD genetic risk in different FTLD subtypes. We used frontal cortex DNA methylation profiles from three FTLD datasets containing different subtypes of FTLD-TDP and FTLD-tau: FTLD1m (N = 23) containing FTLD-TDP *C9orf72* mutation carriers and sporadic cases, FTLD2m (N = 48) containing FTLD-Tau *MAPT* mutation carriers, FTLD-TDP *GRN* and *C9orf72* mutation carriers, and FTLD3m (N = 163) sporadic FTLD-Tau (progressive supranuclear palsy - PSP) cases, and corresponding controls. We then leveraged FTLD transcriptomic and proteomic datasets to investigate possible downstream effects of DNA methylation changes. Our analysis revealed shared promoter region hypomethylation in *STX6* across FTLD-TDP and FTLD-tau subtypes, though the largest effect size was observed in PSP cases compared to controls (delta-beta = −32%, FDR adjusted-*p* value=0.002). We also observed dysregulation of the *STX6* gene and protein expression in some FTLD subtypes. Additionally, we performed a detailed examination of *MAPT*, *GRN* and *C9orf72* across subtypes and observed nominally significant differentially methylated CpGs in variable positions across the genes, often with unique patterns and downstream changes in gene/protein expression in mutation carriers. We highlight aberrant DNA methylation at different CpG sites mapping to genes previously associated with genetic risk of FTLD, including *STX6*. Our findings support convergence of genetic and epigenetic factors towards disruption of risk loci, bringing new insights into the contribution of these mechanisms to FTLD.

## Introduction

Frontotemporal lobar degeneration (FTLD) represents a spectrum of clinically, genetically, and pathologically heterogeneous neurodegenerative disorders characterised by progressive atrophy of the frontal and temporal lobes of the brain (1,2). FTLD is the umbrella term that describes the neuropathology of frontotemporal dementias (FTD) and related disorders. FTD is the second most common form of early-onset dementia and FTD also represents an estimated 25% of dementia cases occurring in individuals over 65 (3,4). Damage to frontal and temporal regions of the brain typically manifests as executive dysfunction, changes in personality and behaviour and language deficits within the clinical subtypes of FTD; behavioural variant frontotemporal dementia (bvFTD), logopenic variant primary progressive aphasia (lvPPA), semantic variant PPA (svPPA)/semantic dementia (SD), nonfluent variant or progressive nonfluent aphasia (PNFA) (5). Amyotrophic lateral sclerosis (ALS) and atypical parkinsonian syndromes, including progressive supranuclear palsy (PSP), frontotemporal dementia and parkinsonism linked to chromosome 17 (FTDP-17) and corticobasal degeneration (CBD), overlap with the clinical phenotypes of FTD and are also neuropathologically classed under the FTLD umbrella (6).

The neuropathological classification of FTLD is based on the presence and morphology of protein aggregates: 50% of cases are attributed to the presence of TAR DNA-binding protein (TDP-43) positive aggregates (FTLD-TDP) (which is further divided A-E subtypes according to the genetic contribution and distribution of the aggregates), 40% to neuronal and glial inclusions of tau (FTLD-tau), while the remaining 10% is comprised of cases with inclusion bodies showing immunoreactivity for fused in sarcoma (FTLD-FUS) and FTLD-UPS involving protein inclusions of the ubiquitin proteasome system in individuals affected by a mutation in *CHMP2B*. A minority of cases show no known proteinaceous inclusions and are classified as FTLD-ni (2).

FTLD is reported to have a strong genetic component, with 30-50% of cases having a positive family history with at least one affected close relative (7). Heritability varies greatly between syndromes, with frequency of mutations also different between geographical populations (8). Most of the heritability in European populations is attributed to autosomal dominant mutations in three genes: Chromosome 9 open reading frame 72 (*C9orf72*), progranulin (*GRN*), and microtubule-associated protein tau (*MAPT*) (9–12). Rare mutations in other genes, including *TARDBP*, *VCP* and *TBK1,* have also been associated with inherited forms of FTLD (13). However, many FTLD cases are sporadic, and several genetic risk factors have been identified through genome-wide association studies (GWAS) (14–18). Single nucleotide polymorphisms (SNPs) in *MAPT* and *MOBP* loci have been associated with risk of FTD and PSP suggesting common genetic denominators across subtypes of FTLD (18–20). SNPs in *STX6* and *EIF2AK3* have been reported to influence the risk of PSP, with no reported association with risk of FTD so far. Exploring the contributions of mutation carriers to the disease phenotype has been an avenue to elucidate which signatures are unique to causative genes (21–25). Although the identification of these FTLD risk genes has provided a basis for exploring pathways and mechanisms driving the pathology of these diseases, genetics on its own has not explained the clinicopathological heterogeneity of FTLD. Epigenetic modifications such as DNA methylation reflect the interplay between genetics and the environment. These modifications are regulatory mechanisms which influence gene expression without changing the underlying DNA sequence. As most human diseases, including neurodegenerative diseases, result from gene deregulation with loss or gain in their functions, epigenetic modifications influencing disease are gaining attention (26–29).

We note that DNA methylation contributes to tight gene expression regulation, as this mechanism has been reported to contribute to changes in expression of the major FTD genes *GRN* and *C9orf72* in FTLD individuals compared to controls (30–33). There has been no conclusive evidence to link DNA methylation at *MAPT* to changes in its expression levels, despite preliminary suggestions of an effect in PSP (30–34). To further assess the relevance of DNA methylation in FTLD, we previously published an epigenome-wide association study (EWAS) meta-analysis using post-mortem frontal lobe DNA methylation profiles from three datasets comprised of different subtypes of FTLD-TDP and FTLD-tau (35). As ageing is a key risk factor for neurodegeneration, we have also investigated biological ageing in FTLD by using DNA methylation clocks (36,37). The results provided more evidence for the involvement of variable DNA methylation in FTLD pathogenesis and accelerated ageing (35,38).

For this study, we compiled a list of causal and risk genes associated with FTLD and leveraged *omics* data from available brain derived datasets. We investigated DNA methylation patterns in the FTLD genetic risk-related loci and determined whether the patterns varied across the heterogeneous FTLD subtypes. As DNA methylation plays a key role in regulating gene expression, we also investigated possible downstream dysregulation in gene and protein expression using transcriptomics and proteomics data. One of our main findings was dysregulation of DNA methylation at the Syntaxin-6 (*STX6*) locus across FTLD-TDP and FTLD-tau subtypes. We also report on DNA methylation patterns and further dysregulation with the major FTLD Mendelian loci (*C9orf72*, *GRN* and *MAPT*). Our findings highlight that loci previously associated with FTLD genetic risk can also be affected via aberrant DNA methylation.

## Methods

### Characterisation of post-mortem brain donors included in DNA methylation investigations

The details of the DNA methylation datasets used in this study are as previously described (35) (Fig. 1, Supplementary Table 1). The post-mortem tissues for FTLD1m (N=23) were obtained from brains donated to the Queen Square Brain Bank where the tissues are stored under a licence from the Human Tissue authority (No. 12198). The brain donation programme and protocols have been granted ethical approval for donation and research by the NRES Committee London Central. The post-mortem tissues for FTLD2m (N=48) were obtained through a Material Transfer Agreement with the Netherlands Brain Bank, as described by Menden *et al*. (39). The data used for FTLD3m (N = 163, after quality control) were made available by Weber *et al*. (40) and accessed through the Gene Expression Omnibus (GEO) database (GEO accession number GSE75704).

**Figure 1.**
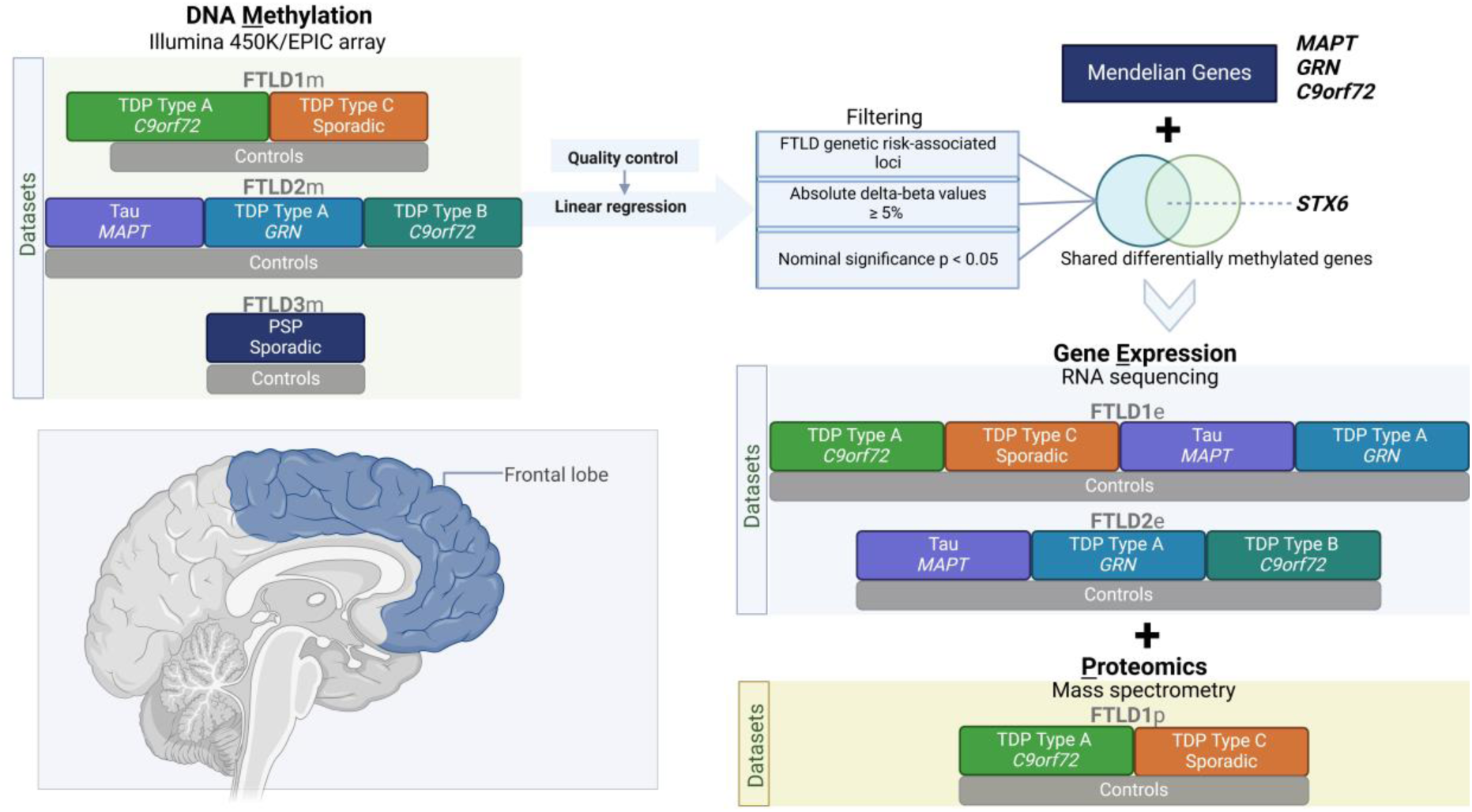
Overview of the study design, datasets and analysis framework. FTLD – Frontotemporal lobar degeneration, PSP – Progressive supranuclear palsy.

### Compilation of known FTLD-associated loci

To focus this study on FTLD genetic risk-associated loci, we compiled a list of genes by searching the DisGeNET text-mining database by disease terms “Frontotemporal lobar degeneration” and “Progressive supranuclear palsy” alongside a literature search to validate entries to the list. Duplicated genes, those that presented with negative results and those where the findings were neither substantial nor replicated were removed. The final list of genes is shown in Supplementary Table 2.

### DNA methylation patterns in FTLD-associated loci

The genome-wide DNA methylation profiles for FTLD1m (N = 23), FTLD2m (N = 48) and FTLD3m (N = 163) were generated using either the Illumina 450K or the EPIC array, as described by Fodder *et al.* (35), Menden *et al.* (39) and Weber *et al.* (40), respectively. Beta-values between 0 and 1 were used to represent the percentage of methylation at each CpG site based on the intensities of the methylated and unmethylated alleles. All analyses and quality control measures were performed using R with Bioconductor packages, as previously described (27,35). Briefly, stringent and harmonised quality control measures were performed on the three datasets through the following steps: 1. the raw data files (idat) were imported for preprocessing, 2. quality control was performed using the minfi (41), wateRmelon (42), and ChAMP (43) packages where cross-reactive probes and probes of poor quality, those mapping to common genetic variants and those mapping to X or Y chromosome, as well as samples with high failure rate (≥ 2% of probes), inappropriate clustering and mismatch of predicted and phenotypic sex, were excluded as previously reported (35). ChAMP Beta-Mixture Quantile (BMIQ) was used to normalise the beta-values which then also underwent logit transformation into M-values for further statistical analysis (44). The annotations of CpG sites mapping to FTLD-associated genes were done based on the Illumina arrays manifest files. Making use of the results from the previously conducted dataset-specific EWAS (35), we characterised in depth the FTLD-associated loci (listed in Supplementary Table 2). Further details regarding regression models used for each EWAS are described by Fodder *et al.* (35) and in Supplementary Table 1. For the current study, we have focused on all methylation sites (CpGs) mapping to FTLD-associated loci showing at least nominally significant DNA methylation changes when comparing FTLD subtypes and controls (unadjusted p<0.05) and an absolute delta-beta of at least 0.05 (i.e., mean difference in DNA methylation levels between cases and controls ≥5%), to ensure the reported differences/effects were biologically relevant and not due to possible technical noise. We analysed the DNA methylation patterns both across datasets and subtypes to also determine if the differential methylation patterns were affected by the presence of certain genetic mutations. We present nominal p-values, unless otherwise specified.

### Gene and protein expression patterns of FTLD-associated loci

To assess whether the expression patterns of genes associated with FTLD risk are in concert with the dysregulation of DNA methylation patterns in FTLD, we used available transcriptomics data for FTLD cases and controls. We have used gene expression data from bulk frontal cortex tissue of FTLD-TDP cases and controls (N = 44) from Hasan et *al.* (45) which has overlapping brain donors with a subset of the FTLD1 DNA methylation dataset, henceforth called FTLD1e. We also used transcriptomic data (N = 44) from the same brain donors as the FTLD2 DNA methylation dataset, called FTLD2e (39). Briefly, RNAseq data for both datasets underwent quality control and processing as previously described (45). The limma package was used to calculate normalisation factors accounting for differences in library sizes (46). Genes with low expression levels were removed i.e. genes where the maximum counts per million (CPM) value across all samples was less than 1. The voom function was used to model the mean-variance relationship and transform the counts data into log2 counts per million (log-CPM) values for linear modelling. A linear model was fitted to the transformed data used to adjust for covariates (Supplementary Table 1). For overlapping brain donors with gene expression and DNA methylation datasets, we also performed DNA methylation-gene expression correlations, using the Pearson correlation coefficient (r) with nominal p-values at a threshold of p<0.05.

Further to gene expression analysis, we looked at the proteomics data for the genes with the FTLD-TDP cases and controls as described in our previous study (35), also with brain donors overlapping with FTLD1 (FTLD1p), where protein levels were quantified using frontal cortex homogenate of frozen post-mortem human brain tissue on control (N = 6), FTLD-TDP type A with *C9orf72* repeat expansion (N = 6), and FTLD-TDP type C (N = 6) cases. Samples were pooled per disease group (three cases per pooled sample) to enable deeper coverage of the proteome with higher fractionation. Fold-changes and standard errors between FTLD-TDP subtypes compared to controls were calculated. As there were only two pooled samples per group, no statistical analysis was performed, but an absolute fold-change >1.5 was considered biologically meaningful. In this proteomics dataset, no data was available for some of the genes we have studied in more detail, including *GRN* and *C9orf72*.

## Results

### DNA methylation is dysregulated in FTLD-associated loci

This study examined in detail loci associated with the genetic risk of FTLD to determine whether these may be affected by changes in DNA methylation, possibly leading to downstream consequences on gene and/or protein expression. We leveraged frontal lobe DNA methylation data from three cohorts composed of multiple FTLD-TDP and FTLD-Tau subtypes, previously studied by Fodder *et al*. (35). The DNA methylation patterns observed for the FTLD risk genes (listed in Supplementary Table 2) showing effect sizes of at least 5% (absolute delta-betas ≥0.05) and nominal p-value <0.05, when comparing FTLD and/or its subtypes with the corresponding controls in each cohort, are shown in Figure 2 and described in Table 1. It is of note that several genes, including *MAPT,* show changes in multiple DNA methylation sites. Although DNA methylation changes are observed in promoter regions represented by CpGs mapping to TSS200 (up to 200 bases upstream of the transcription start site) and TSS1500 (200-1500 bases upstream of the transcription start site), many also occur throughout gene bodies and other regions.

**Figure 2.**
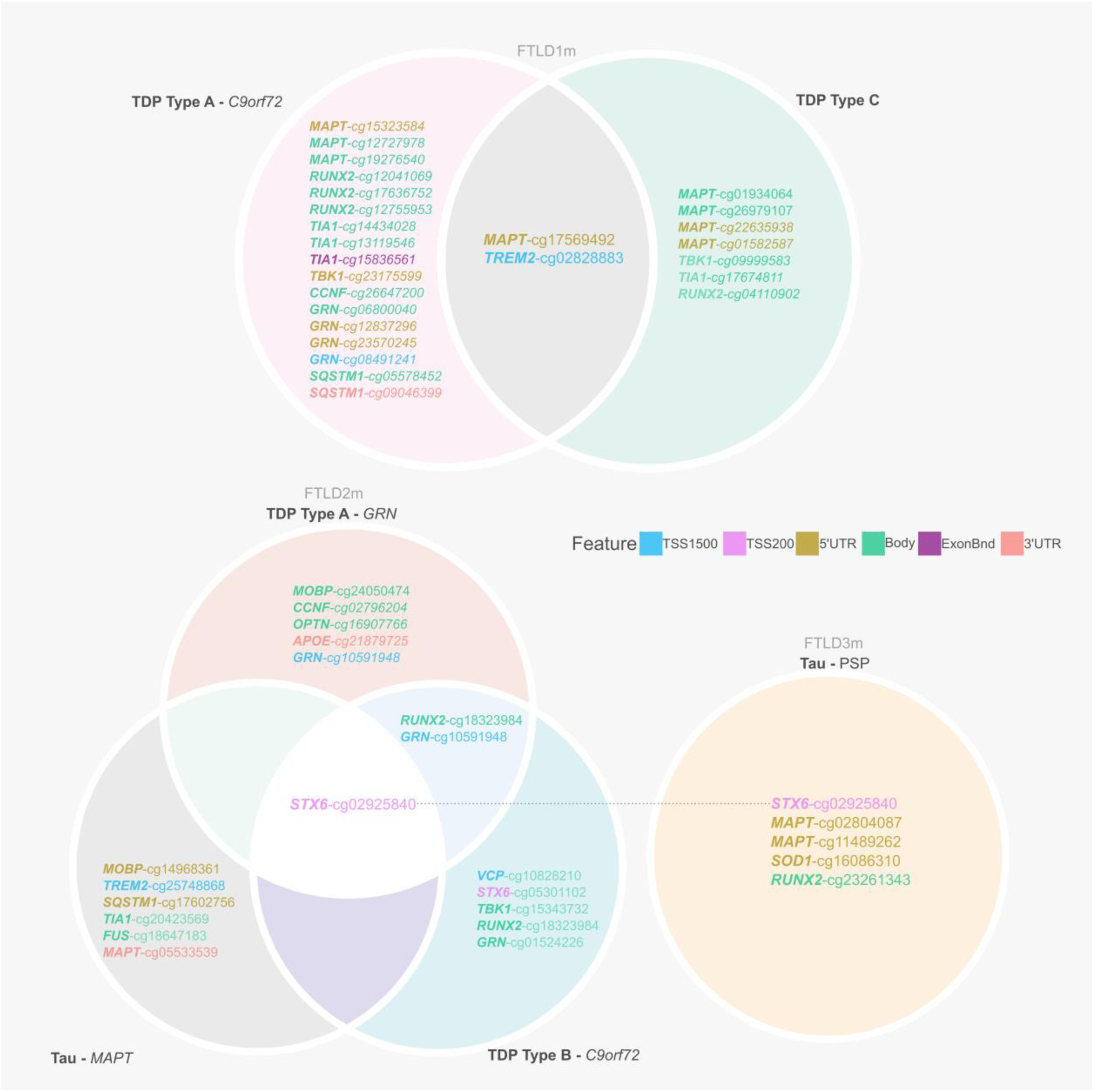
Overview of CpGs showing differences in DNA methylation between FTLD subtypes and controls (absolute delta-beta ≥ 5% and nominal p< 0.05) across three independent datasets (FTLD1m, FTLD2m and FTLD3m). It is of note that dysregulation of cg02925840, mapping to the promoter region of *STX6*, is shared across datasets by all FTLD2m and FTLD3m subtypes. FTLD – Frontotemporal lobar degeneration, PSP – Progressive supranuclear palsy.

**Table 1:** CpGs mapping to FTLD-associated loci showing differential methylation in FTLD cohorts and their subtypes compared to controls (absolute delta-beta ≥ 5% and nominal p< 0.05).

| FTLN1m: FTLN vs CTRL |  |  |  |  |  |  |  |
| --- | --- | --- | --- | --- | --- | --- | --- |
| Gene | CpG | Chr | Position | Feature | CGI | Delta-beta | p-value |
| *MAPT* | cg01934064 | 17 | 44064242 | Body | shelf | -0.14 | 0.024 |
| *MAPT* | cg15323584 | 17 | 44022846 | 5'UTR | shelf | 0.11 | 0.009 |
| *MAPT* | cg17569492 | 17 | 44026659 | 5'UTR | island | 0.09 | 0.019 |
| *MAPT* | cg12727978 | 17 | 44075500 | Body | opensea | 0.08 | 0.009 |
| *TREM2* | cg02828883 | 6 | 41131823 | TSS1500 | opensea | 0.08 | 0.005 |
| *TIA1* | cg14434028 | 2 | 70452453 | Body | opensea | 0.08 | 0.036 |
| *TIA1* | cg13119546 | 2 | 70444039 | Body | opensea | 0.05 | 0.041 |
| *RUNX2* | cg16181497 | 6 | 45409732 | Body | opensea | -0.07 | 0.042 |
| *RUNX2* | cg12755953 | 6 | 45430813 | Body | opensea | 0.06 | 0.039 |
| *RUNX2* | cg04110902 | 6 | 45500999 | Body | opensea | 0.05 | 0.038 |
| *GRN* | cg06800040 | 17 | 42427647 | Body | shelf | 0.07 | 0.022 |
| FTLN1m by subtype: TDP Type A *C9orf72* vs CTRL |  |  |  |  |  |  |  |
| *MAPT* | cg15323584 | 17 | 44022846 | 5'UTR | shelf | 0.17 | 0.002 |
| *MAPT* | cg12727978 | 17 | 44075500 | Body | opensea | 0.15 | 0.001 |
| *MAPT* | cg17569492 | 17 | 44026659 | 5'UTR | island | 0.1 | 0.032 |
| *MAPT* | cg19276540 | 17 | 44060353 | Body | island | 0.08 | 0.035 |
| *RUNX2* | cg12041069 | 6 | 45341222 | Body | shelf | 0.15 | 0.04 |
| *RUNX2* | cg17636752 | 6 | 45391973 | Body | shore | 0.09 | 0.036 |
| *RUNX2* | cg12755953 | 6 | 45430813 | Body | opensea | 0.08 | 0.026 |
| *TIA1* | cg14434028 | 2 | 70452453 | Body | opensea | 0.13 | 0.011 |
| *TIA1* | cg13119546 | 2 | 70444039 | Body | opensea | 0.06 | 0.047 |
| *TIA1* | cg15836561 | 2 | 70442511 | ExonBnd | opensea | 0.06 | 0.028 |
| *TBK1* | cg23175599 | 12 | 64848891 | 5'UTR | shelf | 0.1 | 0.026 |
| *TREM2* | cg02828883 | 6 | 41131823 | TSS1500 | opensea | 0.09 | 0.017 |
| *CCNF* | cg26647200 | 16 | 2482775 | Body | shelf | 0.09 | 0.022 |
| *GRN* | cg06800040 | 17 | 42427647 | Body | shelf | 0.08 | 0.031 |
| *GRN* | cg12837296 | 17 | 42426483 | 5'UTR | opensea | 0.07 | 0.033 |
| *GRN* | cg23570245 | 17 | 42426011 | 5'UTR | opensea | 0.06 | 0.048 |
| *GRN* | cg08491241 | 17 | 42421960 | TSS1500 | opensea | 0.06 | 0.05 |
| *SQSTM1* | cg05578452 | 5 | 179255653 | Body | opensea | 0.07 | 0.005 |
| *SQSTM1* | cg09046399 | 5 | 179264098 | 3'UTR | opensea | 0.06 | 0.025 |
| FTLN1m by subtype: TDP Type C vs CTRL |  |  |  |  |  |  |  |
| *MAPT* | cg01934064 | 17 | 44064242 | Body | shelf | -0.16 | 0.016 |
| *MAPT* | cg17569492 | 17 | 44026659 | 5'UTR | island | 0.08 | 0.045 |
| *MAPT* | cg26979107 | 17 | 44061355 | Body | shore | 0.06 | 0.016 |
| *MAPT* | cg22635938 | 17 | 44039549 | 5'UTR | opensea | -0.06 | 0.012 |
| *MAPT* | cg01582587 | 17 | 44036817 | 5'UTR | opensea | 0.05 | 0.022 |
| *TBK1* | cg09999583 | 12 | 64878162 | Body | opensea | -0.1 | 0.029 |
| *TREM2* | cg02828883 | 6 | 41131823 | TSS1500 | opensea | 0.08 | 0.009 |
| *TIA1* | cg17674811 | 2 | 70443967 | Body | opensea | -0.06 | 0.032 |
| *RUNX2* | cg04110902 | 6 | 45500999 | Body | opensea | 0.06 | 0.023 |
| **FTLD2m: FTLD vs CTRL** |  |  |  |  |  |  |  |
| *TBK1* | cg15343732 | 12 | 64862422 | Body | opensea | 0.09 | 0.034 |
| *STX6* | cg02925840 | 1 | 180992110 | TSS200 | island | -0.08 | 0.003 |
| *MOBP* | cg14968361 | 3 | 39543547 | 5'UTR | shore | 0.08 | 0.050 |
| *TIA1* | cg20423569 | 2 | 70452935 | Body | opensea | -0.05 | 0.008 |
| **FTLD2m by subtype: Tau MAPT vs CTRL** |  |  |  |  |  |  |  |
| *MOBP* | cg14968361 | 3 | 39543547 | 5'UTR | shore | 0.1 | 0.042 |
| *TREM2* | cg25748868 | 6 | 41131213 | TSS1500 | opensea | 0.09 | 0.025 |
| *SQSTM1* | cg17602756 | 5 | 179246001 | 5'UTR | shore | -0.08 | 0.021 |
| *STX6* | cg02925840 | 1 | 180992110 | TSS200 | island | -0.07 | 0.037 |
| *TIA1* | cg20423569 | 2 | 70452935 | Body | opensea | -0.06 | 0.005 |
| *FUS* | cg18647183 | 16 | 31201691 | Body | opensea | 0.06 | 0.042 |
| *MAPT* | cg05533539 | 17 | 44104521 | 3'UTR | opensea | 0.06 | 0.017 |
| **FTLD2m by subtype: TDP Type A GRN vs CTRL** |  |  |  |  |  |  |  |
| *RUNX2* | cg18323984 | 6 | 45386802 | Body | shore | 0.09 | 0.018 |
| *MOBP* | cg24050474 | 3 | 39544326 | Body | shore | 0.09 | 0.046 |
| *CCNF* | cg02796204 | 16 | 2499223 | Body | opensea | 0.08 | 0.031 |
| *STX6* | cg02925840 | 1 | 180992110 | TSS200 | island | -0.08 | 0.015 |
| *OPTN* | cg16907766 | 10 | 13143470 | 5'UTR | shore | 0.07 | 0.041 |
| *APOE* | cg21879725 | 19 | 45412647 | 3'UTR | shore | -0.06 | 0.027 |
| *GRN* | cg10591948 | 17 | 42421375 | TSS1500 | opensea | 0.06 | 0.03 |
| **FTLD2m by subtype: TDP Type B C9orf72 vs CTRL** |  |  |  |  |  |  |  |
| *VCP* | cg10828210 | 9 | 35072977 | TSS1500 | shore | -0.23 | 0.017 |
| *STX6* | cg05301102 | 1 | 180992117 | TSS200 | island | -0.12 | 0.032 |
| *STX6* | cg02925840 | 1 | 180992110 | TSS200 | island | -0.09 | 0.001 |
| *TBK1* | cg15343732 | 12 | 64862422 | Body | opensea | 0.09 | 0.022 |
| *RUNX2* | cg18323984 | 6 | 45386802 | Body | shore | 0.08 | 0.015 |
| *GRN* | cg01524226 | 17 | 42427606 | Body | shelf | 0.06 | 0.007 |
| *GRN* | cg10591948 | 17 | 42421375 | TSS1500 | opensea | 0.06 | 0.03 |
| **FTLD3m: Tau PSP vs CTRL** |  |  |  |  |  |  |  |
| ***STX6*** | **cg02925840** | **1** | **180992110** | **TSS200** | **Island** | **-0.32** | **1.66E-07** |
| *MAPT* | cg02804087 | 17 | 43972969 | 5'UTR | Island | 0.1 | 0.027 |
| *MAPT* | cg11489262 | 17 | 43973426 | 5'UTR | Island | 0.08 | 0.016 |
| *SOD1* | cg16086310 | 21 | 33031992 | 5'UTR | Island | -0.08 | 0.002 |
| *RUNX2* | cg23261343 | 6 | 45413792 | Body | opensea | 0.05 | 0.044 |

**CpGs highlighted in bold** reached epigenome-wide significance (FDR adjusted $p$ -value $\leq 0.05$ ). FTLD – frontotemporal lobar degeneration; PSP – progressive supranuclear palsy; CTRL – controls; CpG – DNA methylation sites; Chr – chromosome; CGI – CpG Islands and other regions; TSS – transcription start site; TSS200 – 0–200 bases upstream of TSS; TSS1500 – 200–1500 bases upstream of TSS; UTR – untranslated region.

### Dysregulation in the *STX6* locus is shared across FTLD subtypes

From our analysis, one CpG mapping to the promoter region of *STX6* (cg02925840) was of particular interest, as it passed genome-wide significance in the FTLD3m dataset (FDR adjusted *p*-value = 0.002), with a strong decrease in methylation levels in the PSP cases compared to controls (delta-beta = −31.5%, Table 1, Fig. 2). Notably, *STX6* has been identified as a genetic risk locus specifically for PSP (20). Still, though to a lesser extent compared to PSP, this same CpG has shown concordant direction of effect in the FTLD2m dataset (FTLD vs controls, delta-beta = −7.9%, nominal p-value = 0.003), and in all its individual subtype comparisons (FTLD-Tau *MAPT* mutants vs controls, FTLD-TDP *C9orf72* mutants and *GRN* mutants vs controls, Table 1). An additional CpG (cg05301102) in the same region showed similar results and reached nominal significance in FTLD2m *C9orf72* mutation carriers and vs controls (delta-beta = −12%, nominal p-value = 0.032) (Fig. 3). Unfortunately, this region could not be analysed in FTLD1m, as probes were excluded during quality control pre-processing of the data (Supplementary Fig. 1). Overall, these findings suggest that disruption of DNA methylation patterns at *STX6* locus might be an important feature shared across FTLD-TDP and FTLD-tau and multiple subtypes, including *MAPT, C9orf72* and *GRN* mutation carriers, in addition to sporadic PSP.

**Figure 3.**
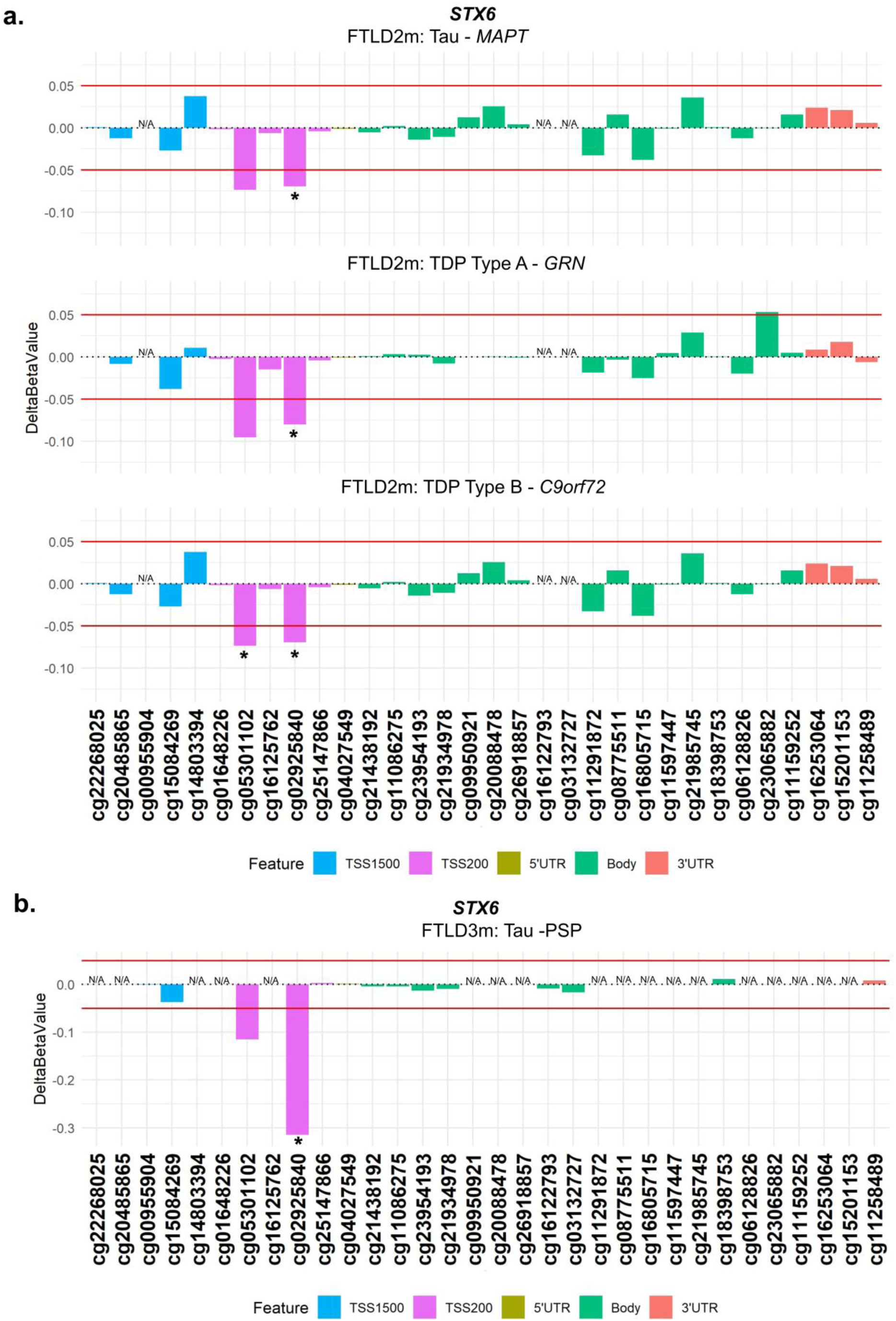
Analysis of DNA methylation patterns across the *STX6* locus reveals that hypomethylation at the promoter region is shared across subtypes in FTLD2m and in FTLD3m (PSP). **(a)** cg02925840 in the promoter region of *STX6* is hypomethylated across subtypes of FTLD2m at the set threshold of an absolute mean difference (delta-beta value) of ≥5% represented by red horizontal lines and at least at nominal significance (*nominal *p* < 0.05). Additionally, cg05301102 also achieves nominal significance in the FTLD-TDP Type A *C9orf72* mutation carriers. **(b)** cg02925840 the promoter region of *STX6* shows strong hypomethylation in FTLD3m in PSP compared to controls (−32%) and reached epigenome-wide significance (FDR adjusted *p* = 0.002). *Indicates nominal p < 0.05. FTLD2m – frontotemporal lobar degeneration DNA methylation cohort 2, FTLD3m – frontotemporal lobar degeneration DNA methylation cohort 3; PSP – progressive supranuclear palsy, TSS – transcription start site; TSS200 – 0–200 bases upstream of TSS; TSS1500 – 200-1500 bases upstream of TSS; UTR – untranslated region. *NA* – These CpGs were not available in the specified dataset due to differences in the methylation array (450K or EPIC) or removal during quality control.

Given this finding in *STX6*, we analysed available FTLD transcriptomics and proteomics datasets to investigate possible downstream consequences in gene and protein expression, respectively. Regarding gene expression (Fig. 4a,b), we observed a small increase in *STX6* expression in the FTLD1e *GRN* mutation carriers (Fold-change 1.2, nominal p < 0.01), only passing multiple testing corrections in FTLD sporadic TDP cases compared to controls (Fold-change 1.1, FDR adj. p = 0.03). We observed, however, a non-significant decrease in *STX6* expression in the *MAPT* and *C9orf72* mutation carriers represented in FTLD2e (Fold-changes −1.1, n.s.). Similarly, gene expression data analysed by Wang et al. (47), showed a non-significant decrease in *STX6* expression in PSP temporal cortex compared to controls. Leveraging a frontal cortex proteomics dataset FTLD1p, we also observed decreased STX6 protein expression in FTLD-TDP type A (*C9orf72* mutation carriers) and FTLD-TDP type C compared to controls (Fold-change −1.9 and −1.5, respectively; Fig. 4c).

**Figure 4.**
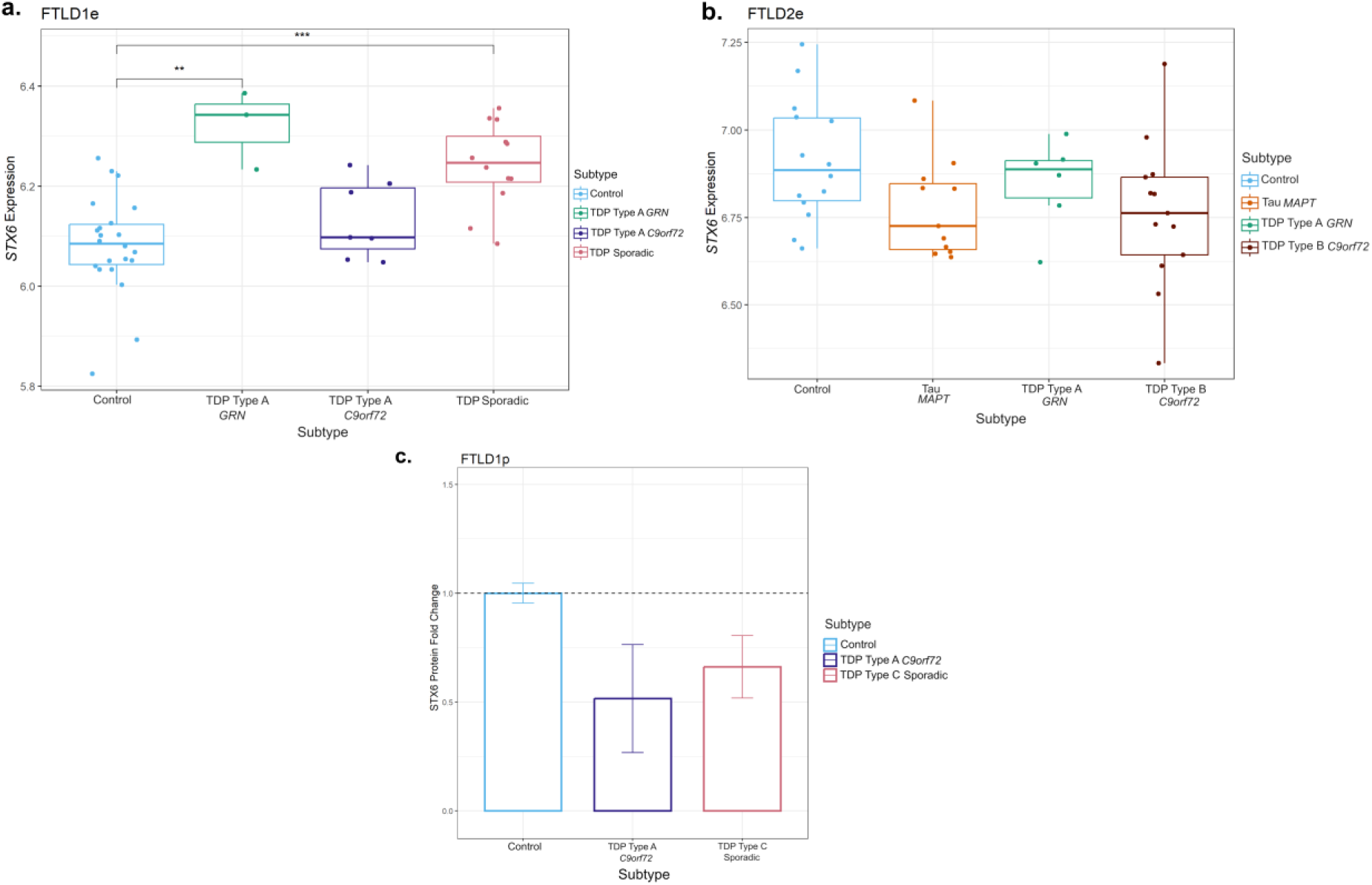
Gene expression and protein patterns of STX6 in frontal cortex of FTLD cases and controls. **(a)** Boxplot showing *STX6* gene expression in FTLD1e cases and controls, with a small increase detected in FTLD-TDP *GRN* mutation carriers and sporadic TDP cases. **(b)** Boxplot of FTLD2e showing *STX6* decreased gene expression but non-significant across all FTLD subtypes (both FTLD-TDP and FTLD-Tau) when compared to controls. Comparisons between the controls and all subtypes for the expression data were carried out using regression models adjusted for multiple covariates as detailed in Supplementary Table 1, nominal p-values are shown (\*\**p* ≤ 0.01; \*\*\**p*≤ 0.001). **(c)** Bar plot showing protein quantifications of STX6 in the frontal cortex of the FTLD1p dataset (FTLD TDP Type A *C9orf72* mutation carriers and FTLD TDP Type C sporadic cases). For each group we used two pooled samples (2 × 3 samples) and derived the quantifications using mass spectrometry. Both FTLD subtypes showed decreased protein expression as visualised by the fold-changes in the bar plot (TDP Type A = −1.9 and TDP Type C = −1.5); standard errors from the mean are also shown. FTLD – frontotemporal lobar degeneration; FTLD1e – gene expression cohort 1; FTLD2e – gene expression cohort 2; FTLD1p – protein expression cohort 1.

Using overlapping cases between FTLD2m and FTLD2e, although non-significant, we observed a positive correlation between *STX6* expression and the top differentially methylated site - cg02925840 in the *MAPT* mutation carriers only (r = 0.42, n.s.; Supplementary Fig. 2), with very weak effects in all other groups. A similar direction of effect was observed for the other highlighted *STX6* CpG - cg05301102 in *MAPT* (r = 0.20, n.s.) as well as in *GRN* mutation carriers (r = 0.63, n.s.). This suggests a possible contribution of DNA methylation shaping *STX6* gene expression landscape at least in these subtypes. However, given the small sample sizes and lack of statistical significance, these findings should be interpreted with caution and warrant further investigation.

### Variable DNA methylation patterns are observed in *MAPT*, *GRN* and *C9orf72*

As mutations in *MAPT*, *GRN* and *C9orf72* represent the majority of familial FTLD cases, we used this opportunity to conduct a detailed investigation of DNA methylation patterns in these loci as well as to analyse possible downstream gene expression in both mutation carriers and non-carriers. Although *C9orf72* did not pass the set thresholds, we still included this locus in our investigation owing to its importance as a Mendelian gene.

#### MAPT

At the *MAPT* locus, which encodes for tau, dysregulation of DNA methylation levels was variable across FTLD datasets and subtypes, not only in FTLD-tau but also in FTLD-TDP, with several CpGs passing the thresholds of absolute delta-beta values ≥5% at nominal significance (\**p* ≤ 0.05, Table 1, Fig. 5, Supplementary Fig. 3). However, the location of those above-threshold CpGs throughout the gene, seemed to differ between FTLD subtypes. The *MAPT* mutation carriers (FTLD-Tau – FTLD2m) had a unique and significantly hypermethylated CpG (cg05533539) in the 3’UTR region of the gene. While the PSP cases (FTLD-Tau – FTLD3m) had two hypermethylated CpGs (cg02804087 and cg11489262) in the 5’UTR region of the gene, none of which surpassed the chosen thresholds or same direction of effect in any other FTLD subtype. For this dataset (FTLD3m), we also stratified samples by the presence/absence of the *MAPT* H2 haplotype to explore whether it could affect DNA methylation patterns at the locus when comparing PSP with controls. However, findings for the gene were similar to those of the unstratified analysis (data not shown), suggesting the *MAPT* haplotypes are not playing a major role in the observed disease-associated DNA methylation landscape at this locus. In the FTLD1m dataset, several CpGs distributed across the gene showed variable methylation patterns, with cg01934064 and cg15323584, in the *MAPT* gene body and 5’UTR, respectively, being the topmost differentially methylated CpGs in FTLD-TDP compared to controls. However, looking at the individual subtypes within this cohort, the cg01934064 was significantly hypomethylated in the TDP type C cases (sporadic) compared to controls, while cg15323584 was hypermethylated in the TDP type A cases (*C9orf72* mutation carriers) compared to controls. The cg17569492 probe, also in the 5’UTR region, was hypermethylated in both FTLD-TDP types A and C. Further to this finding, it is of note that FTLD-TDP types A and C showed downregulation of *MAPT* protein expression compared to controls in the FTLD1p dataset (Fold-changes < −1.5, Fig. 5d). In accordance with this, in the FTLD1e expression dataset, *MAPT* expression was lower in the TDP type A *C9orf72* and sporadic TDP cases (which included TDP type C) when compared to controls (nominal p<0.05), while the FTLD2e dataset showed no significant differences between FTLD subtypes and controls. Wang et al. (47) reports a non-significant increase in *MAPT* expression in PSP temporal cortex compared to controls.

**Figure 5.**
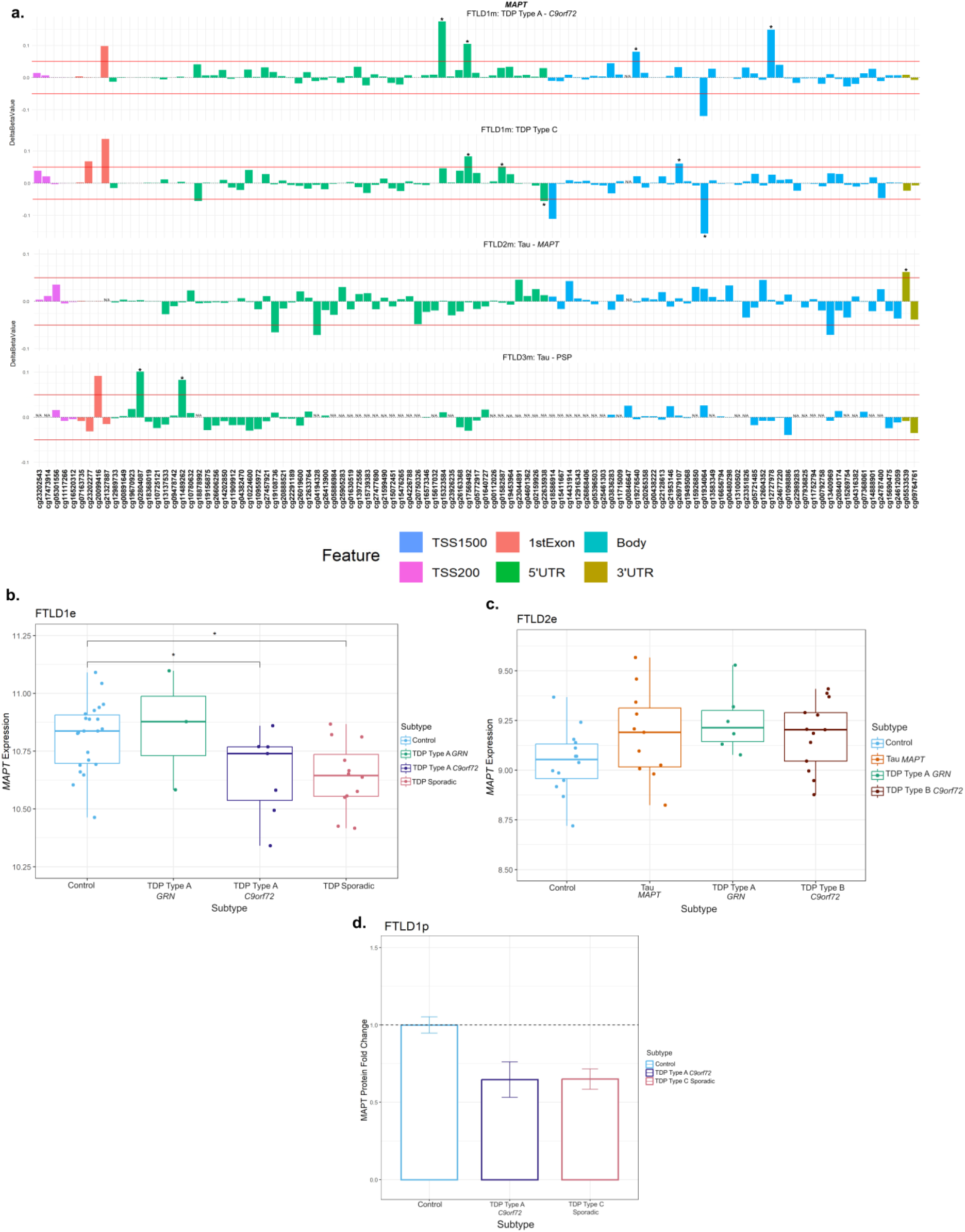
Mixed DNA methylation patterns in the UTRs and Body of *MAPT* with patterns of lower gene and protein expression. **(a)** *MAPT* mutation carriers had one hypermethylated CpG in the 3’UTR region which passed both thresholds of an absolute mean difference of ≥5% and at least at nominal significance (*nominal p < 0.05). Other probes surpassing these thresholds were in the 5’UTR region and the body of the gene. These showed mixed patterns of hypo- and hyper-methylation across different subtypes. **(b)** Boxplot of the FTLD1e cohort showed lower expression of *MAPT* in all subtypes with TDP Type B *C9orf72* and sporadic TDP cases achieving significance. *Indicates *p* ≤ 0.05. **(c)** Boxplot of the FTLD2e cohort showed no changes in *MAPT* expression in the subtypes. Comparisons between the controls and all subtypes for the expression data were carried out using regression models adjusted for multiple covariates as described in Supplementary Table 1. **(d)** Barplot of the FTLD1p cohort where both FTLD subtypes showed decreased protein expression as visualised by the fold-changes (TDP Type A = −1.5 and TDP Type C =-1.5); standard errors from the mean are also shown. FTLD1e – frontotemporal lobar degeneration gene expression cohort 1; FTLD1p – frontotemporal lobar degeneration protein quantification cohort 1. TSS – transcription start site; TSS200 – 0–200 bases upstream of TSS; TSS1500 – 200-1500 bases upstream of TSS; UTR – untranslated region. *NA* – These CpGs were not available in the specified dataset due to differences in the methylation array or removal during quality control.

Using overlapping cases between FTLD2m and FTLD2e, although non-significant, we observed a negative correlation between *MAPT* expression and the 3’UTR CpG which was hypermethylated (cg05533539) in the *MAPT* mutation carriers (r = −0.27, n.s.; Supplementary Fig. 4). It is of note that the controls (r = 0.47, p = 0.089) and the *C9orf72* mutation carriers showed the opposite direction of effect with positive correlations (r = 0.53, p = 0.064). Once again, these findings should be interpreted with caution due to the small sample sizes and lack of statistical significance, and warrant further investigation in future studies.

#### GRN

At the *GRN* locus we observed promoter hypermethylation in the TDP Type A cases containing different mutation carriers (FTLD1 TDP Type A – *C9orf72*, FTLD 2 TDP Type A – *GRN*) compared to corresponding controls, the CpGs cg08491241 and cg10591948, respectively, mapping to the TSS1500 region (Fig. 6a, Supplementary Fig. 5). The TDP Type A *C9orf72* subtype also showed CpGs with hypermethylation patterns in the 5’UTR and body while the TDP Type B *C9orf72* subtype had one CpG in the body surpass the set thresholds (absolute delta-beta ≥5% and nominal p < 0.05). In both the FTLD1e and FTLD2e datasets, when compared to controls, higher expression was observed in the *MAPT* and *C9orf72* mutation carriers while lower expression was observed in the FTLD-TDP Type A *GRN* cases in both datasets (Fold-changes of −1.2 and −1.6, respectively), as often observed with promoter hypermethylation, though this effect only achieved nominal statistical significance in FTLD2e (Fig. 6b-c).

**Figure 6.**
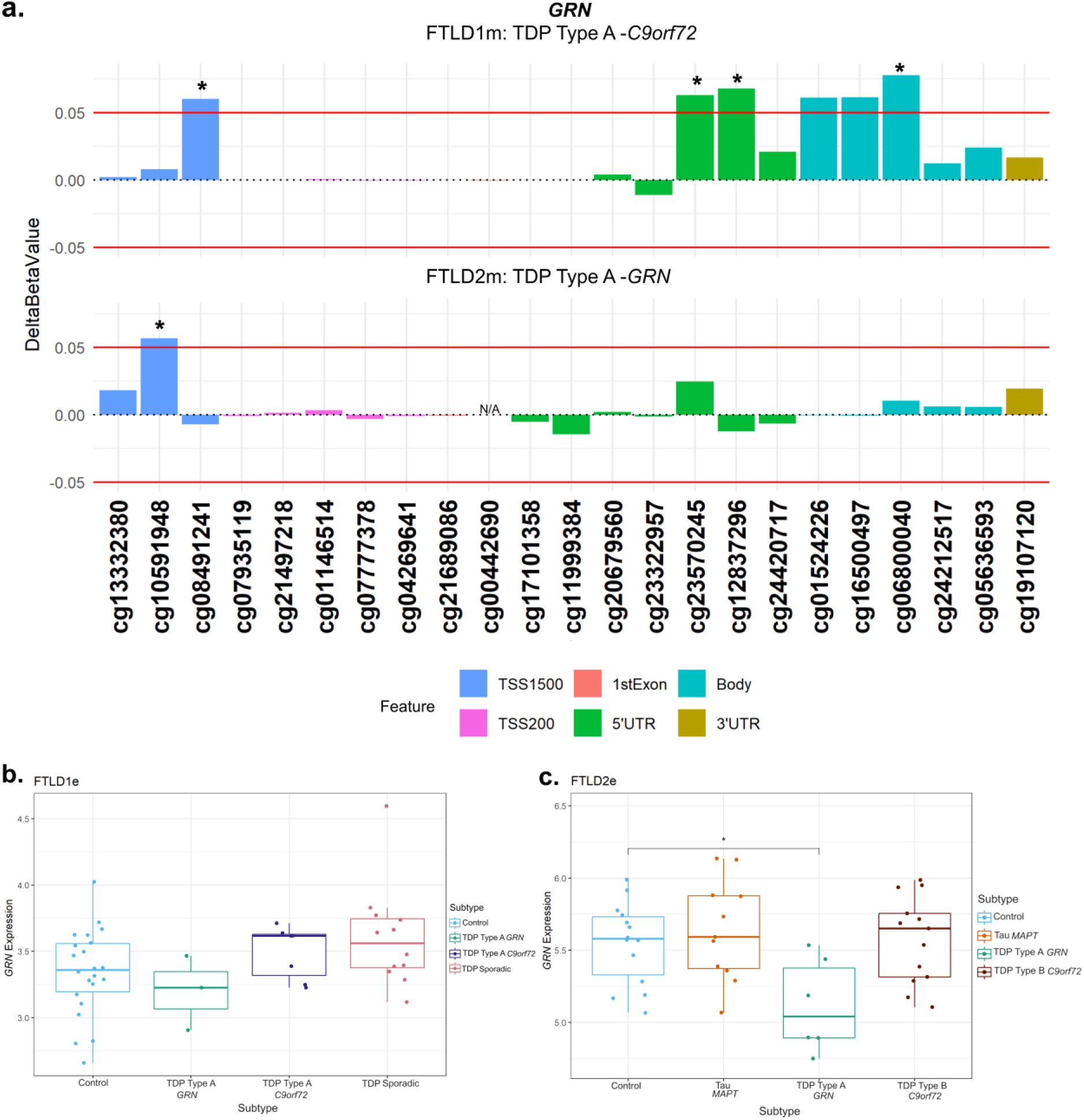
Promoter hypermethylation in *GRN* in FTLD-TDP Type A cases with lower gene expression in *GRN* mutation carriers. **(a)** The TDP Type A cases in both the FTLD1m and FTLD2m cohorts with different mutation carriers (*C9orf72* and *GRN*, respectively) showing hypermethylation in the promoter region of *GRN* passing both thresholds of an absolute mean difference of ≥5% and at least at nominal significance (nominal p < 0.05*). The *C9orf72* carriers also showed above-threshold hypermethylation in CpGs in the 5’UTR and gene body of *GRN.* **(b)** Boxplot of the FTLD1e cohort with mixed expression patterns of *GRN* in FTLD subtypes compared to controls with none achieving statistical significance after multiple testing corrections. **(c)** Boxplot of the FTLD2e cohort showing mixed patterns of gene expression with the FTLD-TDP Type A *GRN* mutation carriers showing a nominally significant decrease in gene expression. Comparisons between the controls and all subtypes for the expression data were carried out using regression models adjusted for multiple covariates as described in Supplementary Table 1. *Indicates nominal p < 0.05. Note: the *GRN* mutation carriers were observed to show lower expression than the corresponding controls and other subtypes in both FTLD1e and FTLD2e. FTLD1e – frontotemporal lobar degeneration gene expression cohort 1; FTLD2e – frontotemporal lobar degeneration gene expression cohort 2; TSS – transcription start site; TSS200 – 0–200 bases upstream of TSS; TSS1500 – 200-1500 bases upstream of TSS; UTR – untranslated region. *NA* – These CpGs were not available in the specified dataset due to differences in the methylation array or removal during quality control.

#### C9orf72

As *C9orf72* is an important gene in FTLD, even though no CpG fully met the established thresholds (absolute delta-beta ≥5% and p-value <0.05), we still detailed the DNA methylation patterns throughout the locus as well as downstream gene expression changes (Fig. 7). We observed higher DNA methylation levels with a delta-beta > 5% (n.s.) in two CpGs only in *C9orf72* mutation carriers (both the TDP Type A and TDP Type B subtypes) compared to controls (Fig. 7a, other subtypes are shown in Supplementary Fig. 4). One near the location of the *C9orf72* repeat expansion in the 5’UTR region and another within the promoter region (cg01861827 and cg14363787, respectively). We also observed a significant downregulation of *C9orf72* gene expression in *C9orf72* mutation carriers, both in FTLD-TDP types A (Fold-change = −1.3, nominal p = 0.005) and B (Fold-change = −1.9, FDR adj. p = 0.003), when compared to the corresponding controls (Fig. 7 b,c). Although to a lesser extent, decreased *C9orf72* expression was also observed in *GRN* and *MAPT* mutations carriers compared to controls, suggesting this locus may be more broadly dysregulated across FTLD subtypes.

**Figure 7.**
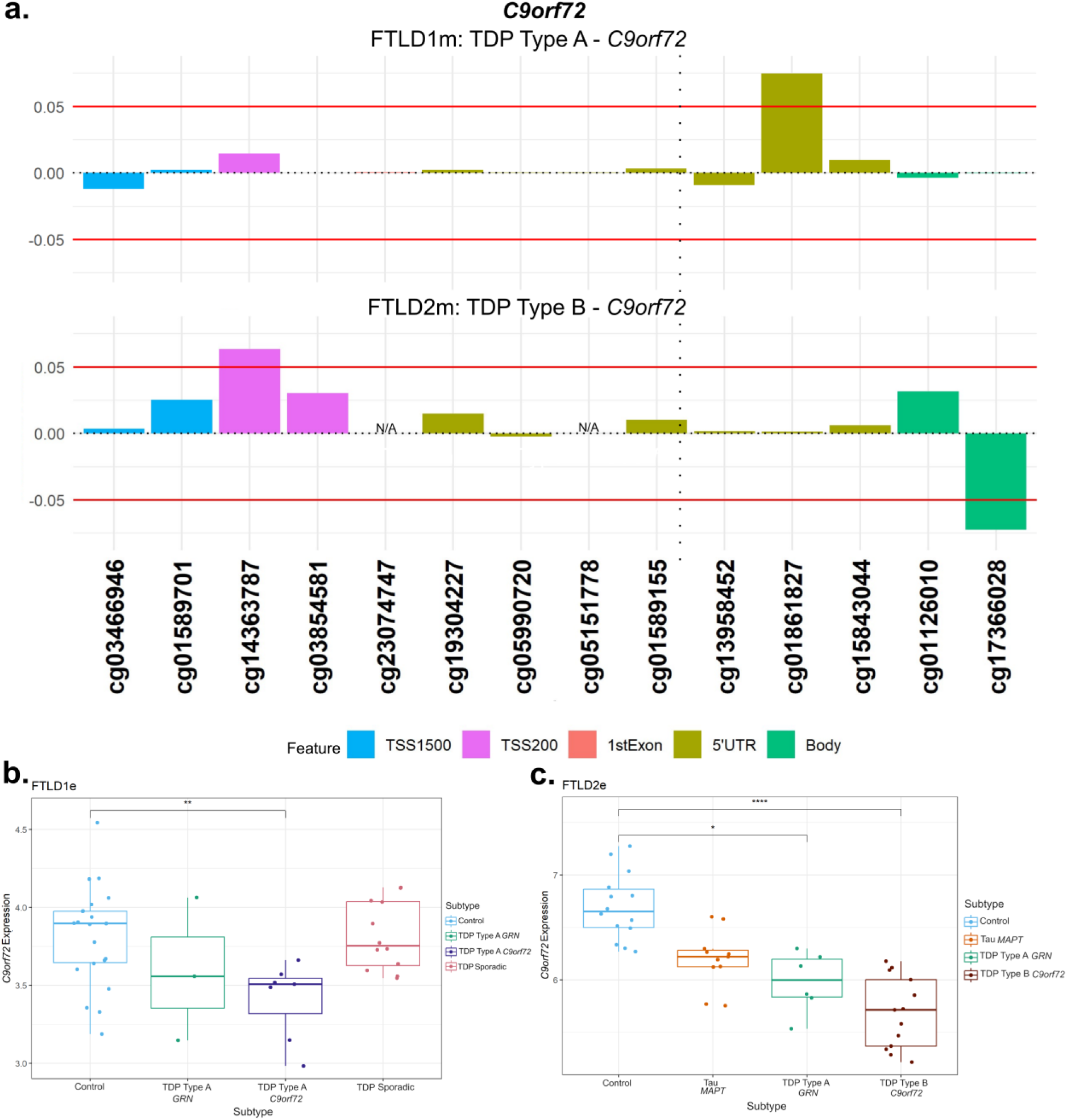
DNA methylation and gene expression patterns in *C9orf72*. **(a)** Only *C9orf72* mutation carriers (TDP Type A and TDP Type B) showed higher levels of methylation compared to the corresponding controls. The TDP Type A cases (FTLD1m) showed this effect in a CpG at the 5’UTR region while the TDP Type B cases (FTLD2m) showed the hypermethylation in a CpG at the promoter region. These CpGs passed the threshold of an absolute mean difference of ≥5% although did not achieve nominal significance (nominal p > 0.05). The vertical dotted line in the 5’UTR region represents the approximate location of the *C9orf72* G_4_C_2_ hexanucleotide repeat expansion. **(b)** Boxplots of the FTLD1e cohort showing lower expression of *C9orf72* all subtypes with the TDP Type A *C9orf72* mutation carriers only achieving nominal statistical significance. ** Indicates nominal *p* ≤ 0.01 **(c)** Boxplots of FTLD2e where all subtypes showed lower gene expression when compared to controls with only TDP Type B *C9orf72* mutation carriers achieving statistical significance after multiple testing corrections and TDP type A *GRN* carriers achieving nominal significance * *p* ≤ 0.05; **** *p* ≤ 0.0001. While all the FTLD cases showed a decrease in expression compared to controls, the *C9orf72* mutation carriers were observed to show the largest effect size. FTLD1e – frontotemporal lobar degeneration gene expression cohort 1; FTLD2e – frontotemporal lobar degeneration gene expression cohort 2; TSS – transcription start site; TSS200 – 0–200 bases upstream of TSS; TSS1500 – 200-1500 bases upstream of TSS; UTR – untranslated region. *NA* – These CpGs were not available in the specified dataset due to differences in the methylation array or removal during quality control.

## Discussion

FTLD has a strong genetic component both in terms of Mendelian genes and in genes associated with risk in sporadic cases. However, genetics alone cannot explain the clinicopathological heterogeneity and/or overlap between FTLD subtypes. This suggests that epigenetic regulatory mechanisms, such as DNA methylation, that represent the interplay between the genetic makeup of an individual and their environmental exposures, may be at play in FTLD. We therefore set up this study to investigate whether DNA methylation changes could contribute to dysregulation of known FTLD genetic risk-associated loci, and how this is affected. For this study, we used DNA methylation data derived from frontal cortex tissue of three independent cohorts, including FTLD-TDP and FTLD-tau pathology, which we had investigated previously from a different perspective (35). We also combined these datasets with overlapping or corresponding gene and protein expression datasets (35,45) to further characterise possible dysregulation of such FTLD-associated loci. Our findings highlighted DNA methylation changes in *STX6*, shared across different FTLD subtypes as a major finding. Furthermore, by characterizing DNA methylation and gene expression in known FTLD Mendelian genes (i.e., *MAPT*, *GRN* and *C9orf72*), we found that dysregulation may occur even in non-mutation carriers. To our knowledge this is the first comprehensive analysis of DNA methylation patterns and characterisation of its possible downstream consequences in FTLD-associated loci, in mutation and non-mutation carriers and in a range of FTLD-TDP and FTLD-tau subtypes.

The “DNA methylation paradox” underscores the complex relationship between DNA methylation and gene expression. Promoter DNA methylation has garnered attention owing to a typically inverse correlation with gene expression (48–53). Similarly, DNA methylation in the 5’ untranslated region (UTR) has been inversely correlated with gene expression, while in the 3′ UTR region a positive correlation has been observed (53–55). Our analysis highlighted two hypomethylated CpGs at the promoter region of *STX6* (cg05301102 and cg02925840) in multiple genetic forms of FTLD (all subtypes of FTLD2m, including *MAPT*, *GRN* and *C9orf72* mutation carriers) and in sporadic PSP (FTLD3m), with a much larger effect size in the latter. Interestingly, genetic variants in *STX6* had been significantly associated with risk of PSP (FTLD-tau) in multiple studies (20,56–58). *STX6* encodes syntaxin 6, which is a soluble N-ethylmaleimide sensitive factor attachment protein receptor (SNARE)-class protein involved in regulation of vesicle membrane fusion (59). Although syntaxin 6 is widely expressed in tissues throughout the body, Bock, Lin and Scheller showed in their seminal work that the brain is among the tissues expressing the highest levels of STX6 protein (60). Dysregulation of STX6 expression has been associated with AD risk and faster cognitive decline potentially relating to neuronal circuitry pathways (61,62). It has also been associated with PSP risk, as more specifically the SNP rs1411478 risk allele has been associated with decreased *STX6* expression levels in the white matter (56). Variants in and around *STX6* have also been associated with risk of the prion disease, specifically sporadic Creutzfeldt-Jakob disease, with a recent study showing that genetic upregulation of both gene and protein expression of syntaxin-6 in the brain is associated with the disease risk (63–65). Dysregulated transport of misfolded proteins from the endoplasmic reticulum to lysosomes has been hypothesized as an underlying mechanism of STX6 (20,56). Recently, an expression quantitative trait loci (eQTL) colocalization has been shown for *STX6* specifically in oligodendrocytes and brain regions associated with PSP pathology (58). DNA methylation has been implicated in the development, differentiation, and maintenance of oligodendrocyte lineage cells where *STX6* is highly expressed (58,66) therefore, its dysregulation is likely playing a role in disease. It is also of note that PSP, shows tau pathology in oligodendrocytes in the form of coiled bodies (67).

Tau is a microtubule-associated protein, encoded by the *MAPT* gene, which becomes abnormally phosphorylated leading to aggregation and formation of intracellular filamentous inclusions, consisting of hyperphosphorylated tau, in several neurodegenerative diseases. These diseases are called tauopathies and include AD as well as several diseases under the FTLD umbrella (FTLD-tau) such as PSP, Pick’s disease, corticobasal degeneration (CBD), argyrophilic grain disease, and frontotemporal dementia with parkinsonism linked to chromosome 17 (FTDP-17), most of which are sporadic with the exception of the latter which is caused by mutations in *MAPT* (such as the *MAPT* mutation carriers in FTLD2m) (2,68). Interestingly, the work by Lee et al. shows a link between syntaxins 6 and 8 and tau, more specifically that they are important in mediating tau secretion through their interaction with the C-terminal tail region of tau (69). Additionally, it has been proposed that pathological TDP-43 is spread between cells in an autophagy-dependent, prion-like manner via extracellular vesicles potentially involving STX6 (70–72). Our findings of *STX6* dysregulation across distinct pathologies supports a broader involvement of *STX6* in both FTLD-TDP and FTLD-tau subtypes.

*MAPT* is one of the main Mendelian genes associated with FTLD where individuals harbour autosomal dominant mutations (9,73), influencing alternative splicing patterns, producing imbalances in tau isoforms, and/or production of more aggregation-prone mutant tau protein (74–76). Common genetic variation in the *MAPT* locus is also associated with risk of FTLD-tau in non-mutation carriers (20,58,77,78). *MAPT* sits within a complex locus (79) with large insertion-deletion polymorphisms in a large region of Chromosome 17q that is in complete linkage disequilibrium, resulting in two major haplotypes, H1 and its inverted counterpart H2, as well as some sub-haplotypes (80,81). H1 is the most common haplotype and is associated with increased risk of sporadic FTLD tauopathies, mainly the four-repeat tauopathies, CBD and PSP (20,79–81), while the H2 haplotype is protective for PSP and CBD and has been associated with familial FTD and increased risk for the three-repeat tauopathy, Pick’s Disease (81–83). Li et al. performed DNA methylation analysis in peripheral blood of FTLD cases, including PSP, and concluded that DNA methylation at the region of the *MAPT* locus may influence the risk of developing tauopathies alongside the H1/H2 haplotypes (84). In studies using brain tissue, *MAPT* DNA methylation patterns have been variable and region-specific as investigated in PSP, AD and Parkinson’s Disease (34,85,86).

Previous studies have reported no significant differences in methylation in FTLD-spectrum cases compared to controls (30,87). However, we observed several differentially methylated CpGs at the *MAPT* gene body and UTRs in our cohorts in both FTLD-TDP and FTLD-tau subtypes. We note that while DNA methylation patterns for different CpGs at the 5’UTR were variable, all those passing the significance thresholds were hypermethylated. The *MAPT* mutation carriers had a significantly hypermethylated CpG in the 3’UTR (cg05533539), which was not observed in any of the other FTLD subtypes. We found increased expression of *MAPT* across the FTLD2e subtypes, none of which were statistically significant. On the other hand, we found decreased expression in FTLD1e type A with *C9orf72* mutation cases and in sporadic cases (including TDP type C), consistent with FTLD1p decrease in protein expression. Untranslated regions have roles in regulating gene expression (54,88,89). However, the effect of DNA methylation at these regions remains unclear. *MAPT* has a core promoter around its first exon, but it has also been suggested to have alternative promoters at different transcription start sites (76,90), which also affect the length of 3’ and 5’ UTRs (76,88). Taken together, the complexity of *MAPT*’s structure aligns with its high variability in methylation and gene and protein expression in FTLD. Whether UTRs play a significant role in regulating expression in *MAPT* remains a point for future investigation. Overall, these findings also suggest that the dysregulation at the *MAPT* locus is not confined to *MAPT* mutation carriers or tau pathology, but also extends to non-mutation carriers and those with other FTLD pathologies such as FTLD-TDP.

Mutations in *GRN,* which encodes progranulin, are another major cause of autosomal dominant FTLD. These mutations result in decreased expression and loss-of-function of the mutant allele of *GRN* resulting in haploinsufficiency and reduced expression of progranulin (10,91–93). This is particularly important in a disease context as progranulin is proposed to localise near endosomes and lysosomes to participate in endocytosis, secretion and other related key functions (94–96). Additionally, progranulin is involved in neuroinflammation, axonal growth, development and acts as a neurotrophic factor promoting neuronal survival (97,98). *GRN* has also been suggested as a modifier of risk for sporadic cases of FTLD. However, this finding may be related to a disruption of lysosomal activities chaperoned by *GRN* and requires further investigation (99,100). Still, the proposed role for *GRN* across FTLD in conjunction with an appearance of asymmetric cortical atrophy specific to the mutation carriers has provided a strong argument to determine regulatory mechanisms, including epigenetic mechanisms, influencing *GRN* expression (101,102). Hypermethylation at the promoter region of *GRN* has been inversely correlated with gene expression and therefore reduced *GRN* expression in FTLD in sporadic cases (32,33). Banzhaf-Strathmann *et al.* showed hypermethylation at the promoter region in *GRN* in FTLD compared to AD and PD (33). Likewise, we have shown hypermethylation in the promoter region of *GRN* in FTLD-TDP Type A cases not only in *GRN* but also in *C9orf72* mutation carriers. We observed lower *GRN* expression in the *GRN* mutation carriers only though, possibly emphasizing the impact of genetic variation and suggesting that DNA methylation dysregulation beyond the promoter region, as seen in the *C9orf72* mutation carriers, may act differently and/or in concert with other mechanisms to regulate *GRN* gene expression.

Expansion of the non-coding G_4_C_2_ hexanucleotide repeat in the 5’UTR region of *C9orf72* is the most common cause of familial FTLD (8). The mechanism by which this mutation causes disease remains an area of intense research as multiple pathways have been implicated (103). Like *GRN,* haploinsufficiency with reduced *C9orf72* expression and loss-of-function has been proposed (104). Toxic gain-of-function mechanisms have also been suggested (31,105–110). The biological role of *C9orf72* also remains unclear. However, recent studies observing protein-protein interactions suggest involvement in lysosomal activity, vesicle trafficking, axon growth, regulation of mTORC1 signalling and of inflammation (111,112).

Reduced expression of *C9orf72* has been observed in some mutation carriers. Therefore, DNA methylation patterns have been previously analysed to determine whether there was a role for this reversible mechanism in regulating *C9orf72* expression in FTLD (11,31,113,114). Hypermethylation at *C9orf72* has been observed uniquely in mutation carriers in a region upstream of the repeat (115). As *C9orf72* promoter hypermethylation results in reduced *C9orf72* expression, it is suggested to be a protective mechanism acting against the toxic gain-of-function mechanisms including reducing the amount of RNA foci while also validating a loss-of-function mechanism (115). Our results showed promoter hypermethylation in FTLD-TDP type B *C9orf72* mutation carriers. *C9orf72* expression was reduced in all studied FTLD subtypes but with the largest effect size being observed in the *C9orf72* mutation carriers, both in FTLD1e and FTLD2e. This is in line with a previous study that showed reduced expression of *C9orf72* in repeat expansion mutation carriers as well as *MAPT* and *GRN* mutation carriers, and proposed that additional mechanisms independent of promoter hypermethylation, which is primarily observed in *C9orf72* mutation carriers, regulates *C9orf72* expression across FTLD subtypes (113).

As with other studies, there are several limitations. We examined patterns in DNA methylation between subtypes of FTLD, however, this meant using relatively small sample sizes to compare across subtypes which reduced the statistical power to detect additional genome-wide changes. The available DNA methylation profiles were derived using Illumina 450K/EPIC arrays, which are not comprehensive despite their coverage throughout the genome. This is particularly important for complex genes where not all regions overlap with predefined regions covered in the arrays. As DNA methylation and gene expression may vary depending on the cellular composition and properties of a sample, there may still be differences in the tissue once chipped, which may influence findings in this type of study to some degree and cannot be fully accounted for using statistical approaches. We note that even in those donors that have overlapping samples there may be some sample variability between different omics modalities. We were also limited by the lack of full overlap between samples used to generate DNA methylation and gene expression datasets to further dissect possible downstream consequences. Additionally, we cannot completely exclude the possibility that unmeasured genetic variants, including large structural variants, may have an effect on the detection of DNA methylation changes. However, leveraging available DNA methylomics, transcriptomics and proteomics datasets, we strived to report the most consistent findings, with a meaningful biological effect (e.g., absolute delta-beta ≥5% in group comparisons), and analysed the concordance with previously published studies whenever possible.

In summary, this study explored for the first time a cross-subtype analysis of the contribution of DNA methylation to the dysregulation of FTLD genetic risk loci, with or without the presence of genetic mutations in Mendelian FTLD genes. We highlight *STX6* that showed consistent hypomethylation of the promoter region across FTLD subtypes and cohorts. On that basis, our findings support a role for *STX6* in other FTLD subtypes beyond PSP. We suggest that DNA methylation may be influencing *STX6* gene expression levels, at least in some FTLD subtypes. However, it would be important to replicate and analyse this point further in future studies with larger samples sizes per subtype. Taking into consideration a role in regulation of protein localization and its complex relationship with tau, and possibly TDP-43, the role of syntaxin-6 in various subtypes of FTLD warrants further investigation. Additionally, we focused on the Mendelian genes *MAPT*, *GRN,* and *C9orf72* where we describe patterns of DNA methylation and gene expression and showed that dysregulation is not necessarily unique to mutation carriers. Understanding the mechanisms underlying the dysregulation of such genes, including DNA methylation changes, will be key to the development of therapies. Overall, our findings have shown DNA methylation changes in FTLD-associated genes across FTLD subtypes both in carriers of known genetic mutations and in sporadic cases. We highlight that such epigenetic modifications may be a shared mechanism across FTLD subtypes possibly contributing to the dysregulation of gene expression and can provide new insights into genes associated with disease.

## Ethics approval and consent to participate

The post-mortem tissues used to generate the FTLD1 datasets were obtained from brains donated to the Queen Square Brain Bank, where the tissues are stored under a licence from the Human Tissue authority (No. 12198). The brain donation programme and protocols have been granted ethical approval for donation and research by the NRES Committee London Central. The post-mortem tissues used to generate the FTLD2 datasets were obtained from The Netherlands Brain Bank (NBB), Netherlands Institute for Neuroscience, Amsterdam. All NBB Material has been collected from donors for whom a written informed consent was obtained for a brain donation and the use of the material and clinical information for research purposes. The FTLD3 DNA methylation dataset was accessed through a public repository (GEO accession number GSE75704). As described by Weber *et al*. (40), the use of human brain tissue to generate that dataset had been approved by the ethics committees of the University of Giessen and of the Technical University of Munich. All brain samples were from patients who had given informed consent before death.

## Consent for publication

Not applicable.

## Availability of data and materials

Raw DNA methylation and/or RNAseq data from cohorts FTLD1, FTLD2 and FTLD3 can be accessed via the European Genome-Phenome Archive (accession number EGAD00010002055, EGAD00001008014) and NCBI GEO database (accession number GSE75704, and GSE153960). Additional data is available from the corresponding author upon reasonable request.

## Authors’ contributions

NR contributed to data analysis, prepared the figures, and drafted the manuscript; KF and MM contributed to data analysis and interpretation; CT, RdS, PH, JH, TR, TL contributed with datasets and/or interpretation; CB conceptualised and supervised the study; All authors critically reviewed and approved the manuscript.

## Acknowledgements

The authors would like to thank UCL Genomics centre for processing the EPIC arrays for the for the FTLD1m cohort. The authors would also like to acknowledge the Queen Square Brain Bank (London, UK), and the Dutch Brain Bank, Netherlands Institute for Neuroscience (Amsterdam, Netherlands) for providing brain tissues from FTLD cases and controls. The Queen Square Brain Bank is supported by the Reta Lila Weston Institute of Neurological Studies, UCL Queen Square Institute of Neurology.

