## Supplementary Fig for "DNA methylation as a contributor to dysregulation of *STX6* and other frontotemporal lobar degeneration genetic risk-associated loci"

1 Wakefield Street
London WC1N 1PJ
United Kingdom

**Supplementary Figures**

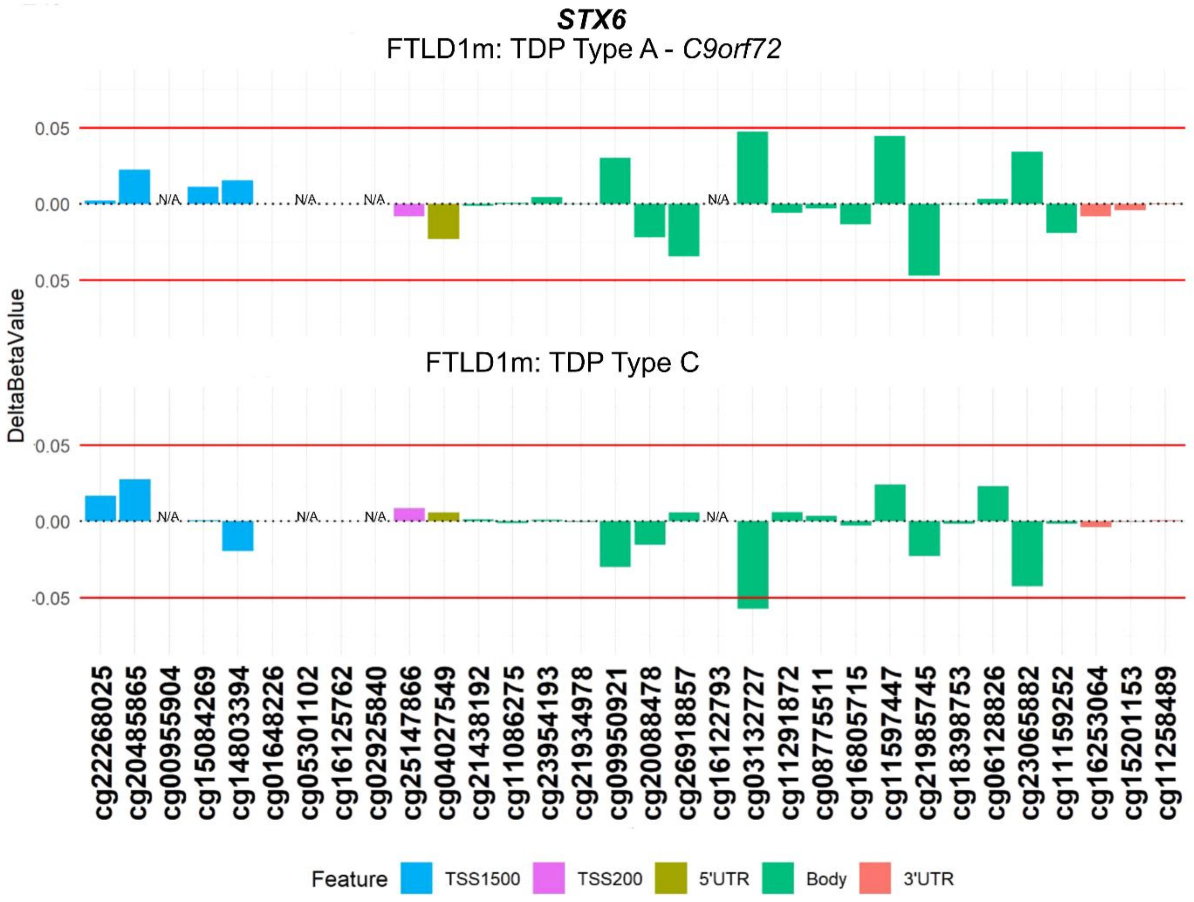


**Supplementary Fig. 1 Analysis of DNA methylation patterns across the *STX6* locus in FTLD1m**. Note: CpGs at the promoter region showing changes in other FTLD subtypes (cg02925840 and cg05301102) were not present in this dataset due to their exclusion during data quality control pre-processing. FTLD1m – frontotemporal lobar degeneration DNA methylation cohort 1, TSS – transcription start site; TSS200 – 0–200 bases upstream of TSS; TSS1500 – 200-1500 bases upstream of TSS; UTR – untranslated region. NA – These CpGs were not available in the specified dataset due to differences in the methylation array (450K or EPIC) or removal during quality control.

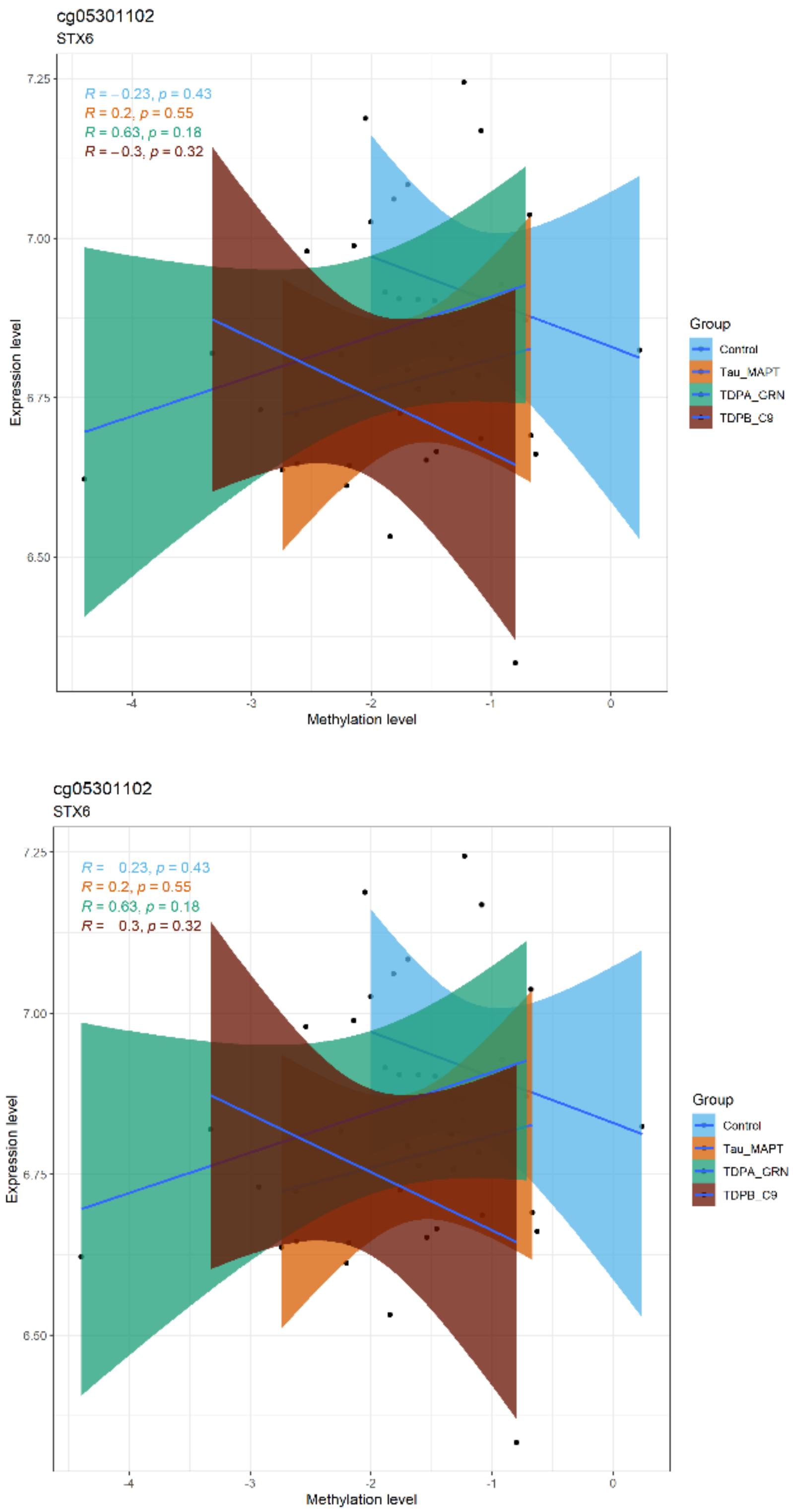


**Supplementary Fig. 2 DNA methylation-gene expression correlations in FTLD2 datasets for the top two CpGs (cg02925840 and cg05301102) mapping to *STX6* promoter**. Log2-transformed gene expression data is shown in the *y*-axis, and DNA methylation levels (M-values) are shown in the *x*-axis. FTLD2 – frontotemporal lobar degeneration cohort 2.

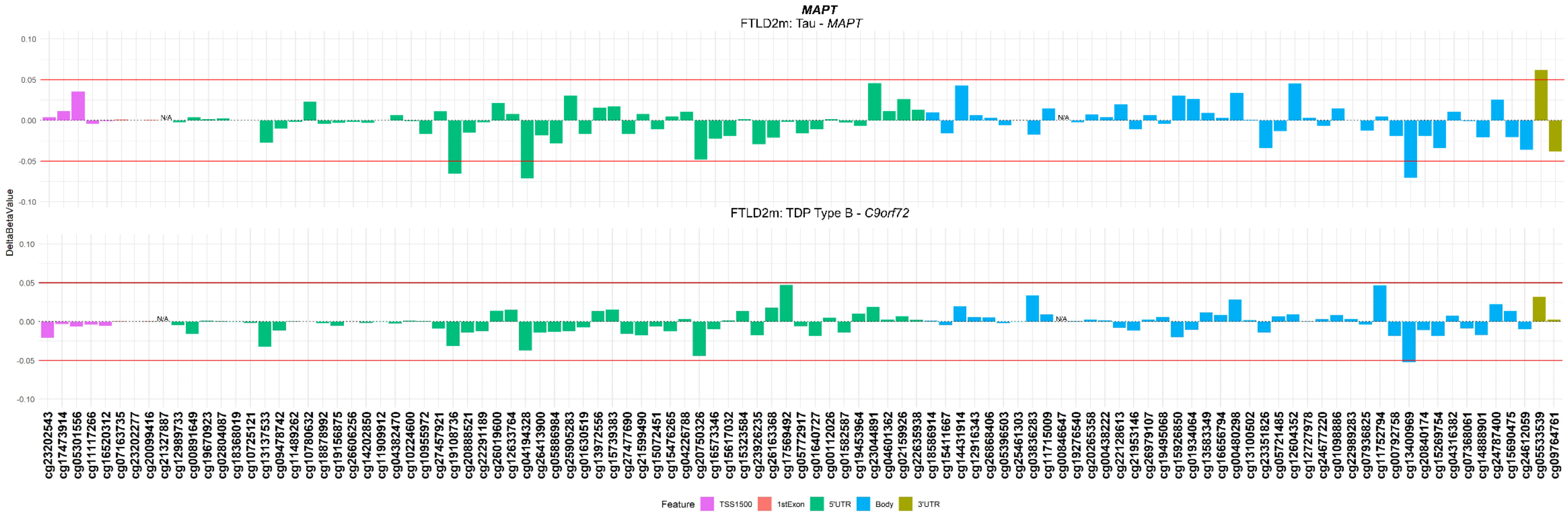


**Supplementary Fig. 3 Analysis of DNA methylation patterns across the *MAPT* locus in *GRN* and *C9orf72* cases from the FTLD2m cohort**. Note: No probes met the significance threshold ($p<0.05$) in the FTLD-TDP Type A *GRN* or Type B *C9orf72* cases compared to controls. FTLD2m – frontotemporal lobar degeneration DNA methylation cohort 2, TSS – transcription start site; TSS200 – 0–200 bases upstream of TSS; TSS1500 – 200-1500 bases upstream of TSS; UTR – untranslated region. NA – These CpGs were not available in the specified dataset due to differences in the methylation array (450K or EPIC) or removal during quality control.

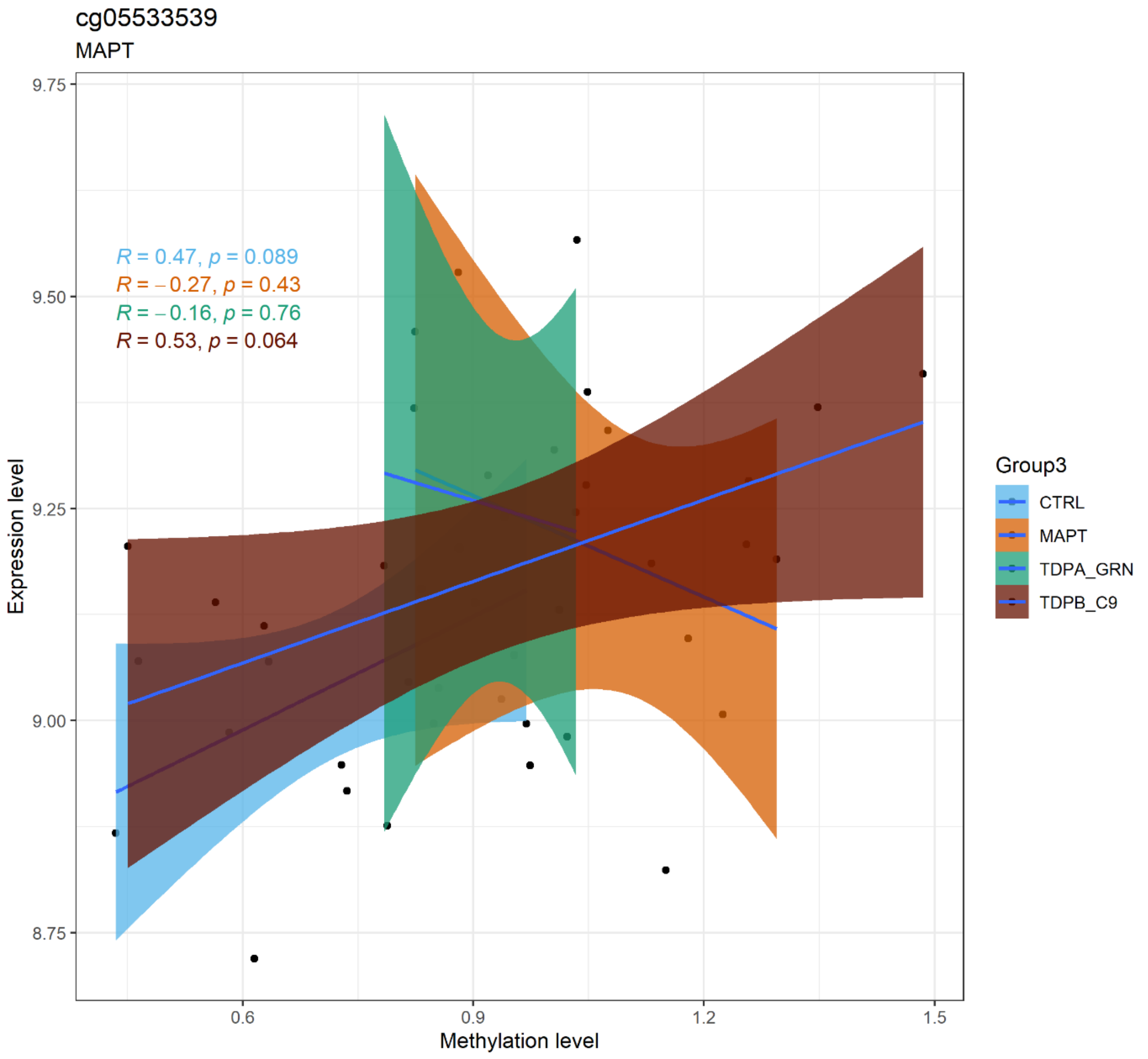


**Supplementary Fig. 4 DNA methylation-gene expression correlations in the FTLD2 datasets for the *MAPT* 3'UTR CpG differentially methylated in *MAPT* mutation carriers (cg05533539).** Log2-transformed gene expression data is shown in the *y*-axis, and DNA methylation levels (M-values) are shown in the *x*-axis. FTLD2 – frontotemporal lobar degeneration cohort 2; 3'UTR – 3' untranslated region.

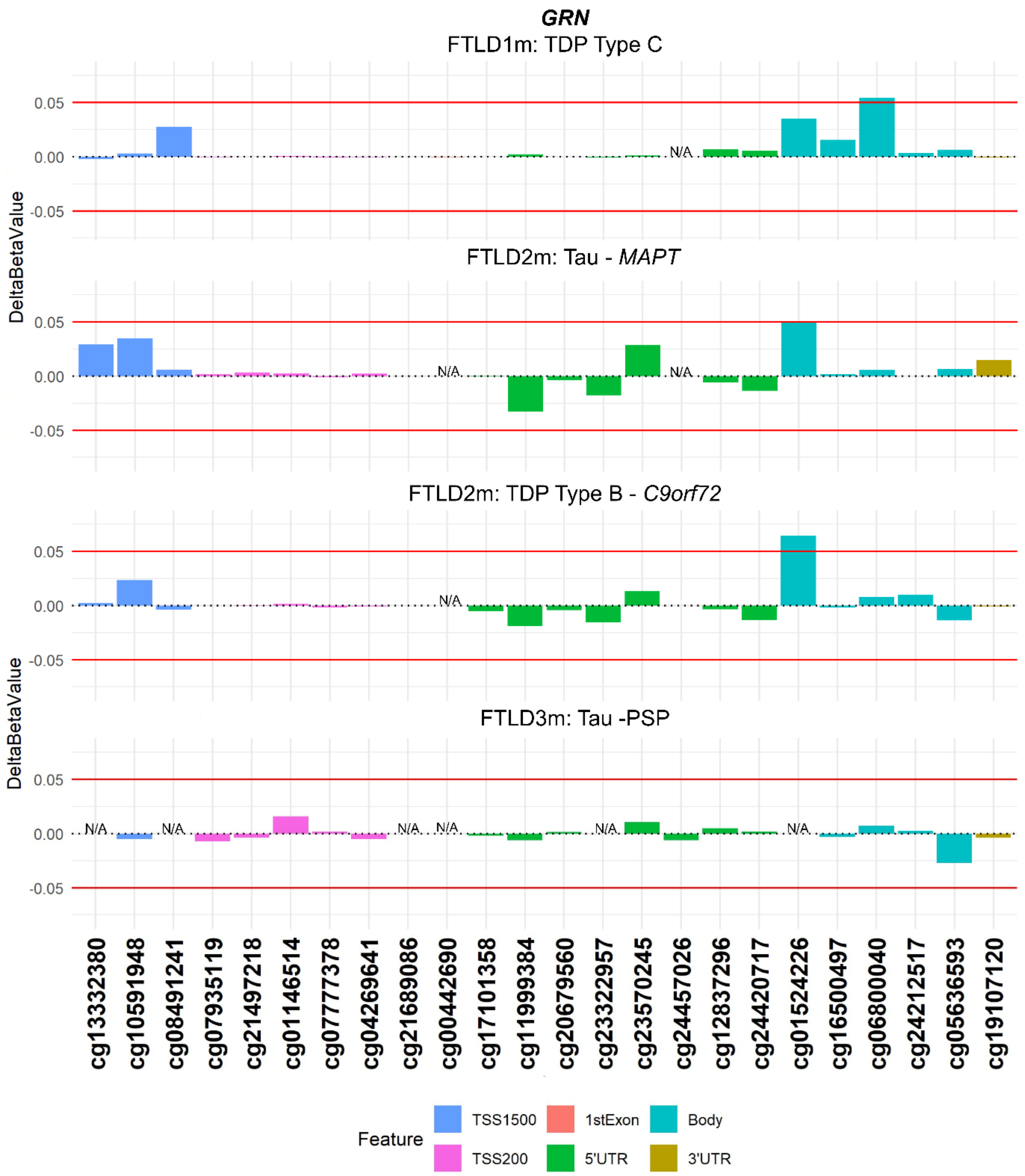


**Supplementary Fig. 5 Analysis of DNA methylation patterns across the *GRN* locus in non-TDP Type A cases**. Note: No probes showed dysregulated methylation (absolute delta-beta ≥5%, p<0.05) at the promoter region in other FTLD-TDP types or FTLD-Tau cases. TSS – transcription start site; TSS200 – 0–200 bases upstream of TSS; TSS1500 – 200-1500 bases upstream of TSS; UTR – untranslated region. NA – These CpGs were not available in the specified dataset due to differences in the methylation array (450K or EPIC) or removal during quality control.

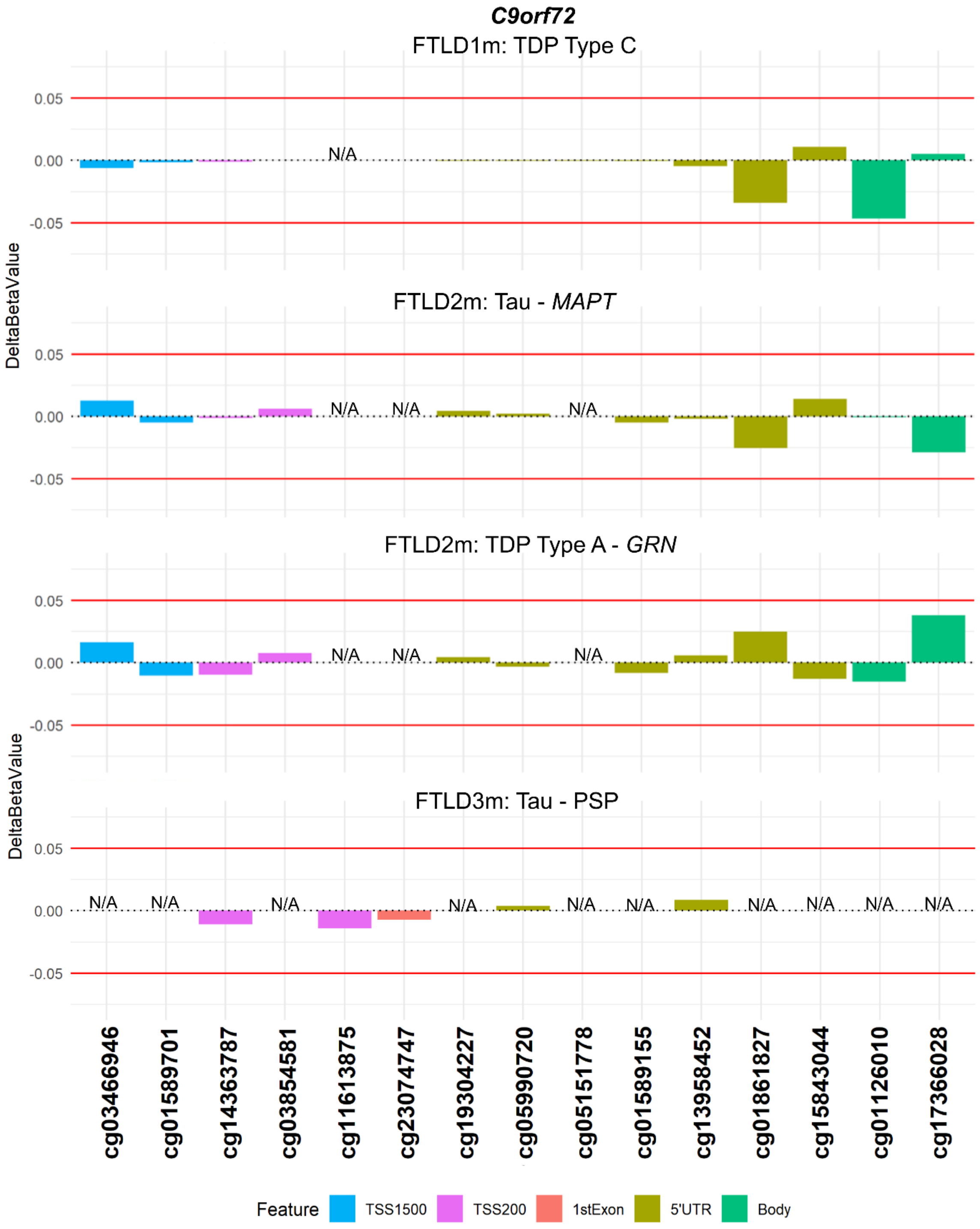


**Supplementary Fig. 6 Analysis of DNA methylation patterns across the *C9orf72* locus in non-mutation carriers**. Note: No probes at any region showed an absolute delta-beta, i.e. mean difference when compared to controls, of ≥5%. TSS – transcription start site; TSS200 – 0–200 bases upstream of TSS; TSS1500 – 200-1500 bases upstream of TSS; UTR – untranslated region. NA – These CpGs were not available in the specified dataset due to differences in the methylation array (450K or EPIC) or removal during quality control.
